# DNA barcodes for rapid, whole genome, single-molecule analyses

**DOI:** 10.1101/450809

**Authors:** Nathaniel Wand, Darren A. Smith, Andrew Wilkinson, Ashleigh Rushton, Stephen J. W. Busby, Iain Styles, Robert K. Neely

## Abstract

We report an approach for visualizing DNA sequence and using these ‘DNA barcodes’ to search complex mixtures of genomic material for DNA molecules of interest. We demonstrate three applications of this methodology; by identifying specific molecules of interest from a dataset containing gigabasepairs of genome; by straightforward strain-typing of bacteria from such a dataset and, finally, by locating infecting virus molecules in a background of human genomic material. DNA barcodes enable quantitative understanding of complex genome mixtures, on a large scale. As a result of the dense fluorescent labelling of the DNA, individual barcodes of the order 40 kilobase pairs in length can be reliably identified. This means DNA can be prepared for imaging using standard handling and purification techniques. The recorded dataset provides stable physical and electronic records of the total genomic content of a sample that can be readily searched for a molecule or region of interest.

## INTRODUCTION

Direct visualization of the DNA sequence by optical mapping offers a unique perspective on genome structure (1). The single-molecule, long-range information that is derived from mapping has recently been invaluable in revealing hundreds of large genomic rearrangements of the human (2, 3) and great ape (4) genomes. However, the development of nanopore sequencing has enabled sequence read lengths that are of the same scale as optical maps (5), thereby providing an alternative approach for scaffolding or assembling sequencing data from a dataset which is far more information rich than the comparative mapping data.

Whilst nanopore sequencing is increasing the competition in the space traditionally occupied by optical mapping, the commercial Bionano Genomics platform continues to offer a valuable and reliable means of deriving a reference scaffold for genomic assemblies (6). Furthermore, imaging is particularly suited to multiplexed experiments, meaning that mapping offers a route by which whole genome studies can be performed, where sequence data can be directly correlated to, for example, a DNA repair event (7), DNA replication (8), or protein binding (9), at the single-molecule level. Whilst the promise of single-molecule mapping approaches for imaging DNA-based events has been demonstrated in such experiments, none to date has approached achieving this on the scale of a whole bacterial or human genome.

Several approaches exist for producing optical maps of DNA sequence (10). Until recently, mapping has been reliant on either restriction enzymes (11) or nicking enzymes (2) to define the sequence motifs used in mapping. However, the discovery that methyltransferase enzymes can be used for fluorescent labelling of DNA (9, 12–14) has been transformative because of their ability to yield sequence-specifically modified DNA without introducing damage (cuts or nicks) to the DNA. Previous mapping approaches have focussed on mapping of long DNA molecules, typically greater than 200 kbp in length, driven by the relative infrequency of mapping sites. Whether defined by restriction, nicking or methyltransferase enzyme, a map must carry enough information for it to be reliably matched to a sequence of interest, and hence, applied. For a 5- or 6-base targeting enzyme, map density is typically one site every 10 kbp, resulting in the need for molecules hundreds of kbp in length for reliable map assembly. The production and handling of DNA molecules on this size range is technically challenging, requiring careful extraction and clean-up of DNA using gels to minimise shearing forces in the sample.

We describe a methodology that significantly increases the accessibility and range of potential applications for optical mapping. We show that a high-density, yet sequence-specific labelling pattern, directed by a DNA methyltransferase allows single DNA molecules to be probed for sequences of interest at the whole-genome scale. Map analysis is demonstrated using relatively small DNA fragments (∼30 kbasepairs) that can be readily prepared using standard DNA extraction kits and protocols. We show how this can be used to build datasets of hundreds of thousands of DNA molecules (several gigabasepairs of DNA) in less than an hour using standard fluorescence microscopy. To extend the imaging to applications in genomics, we have developed image-matching and classification techniques, which enable unique, whole genome analyses to be performed. For example, hundreds-of-thousands of single-molecule barcodes can be queried for a sequence of interest, and barcode images can be clustered, based on similarity, allowing the prevalent DNA molecules in a mixed population to be identified. We demonstrate application to the detection of viral infection in human cells, to strain-typing of bacteria, and to the visualization of a region of interest in a genome that had been modified using CRISPR-Cas9 genome editing. In all, we expect this approach to find widespread application in the quantitative study of mixed genome samples and, particularly in studying the sequence context of genome-wide events and processes, such as replication or protein binding, that are not directly accessible with sequencing technologies.

## MATERIALS AND METHODS

### Labelling of genomic DNA

A 200 μl solution containing 1x CutSmart Buffer (NEB), 10 μg genomic DNA, 0.9 μg TaqI DNA methyltransferase (M.TaqI) and 750 μM AdoHcy-azide (Figure S1) was prepared and incubated at 50°C for 1 hour. Subsequently, 5μl 18mg/ml proteinase K (NEB)/0.1% Triton X-100 (Sigma-Aldrich) was added and this was incubated at 50°C for 1 hour, before purification by GenElute Bacterial Genomic DNA kit (Sigma-Aldrich). DNA was eluted into 200 μl TE Buffer (10 mM tris, 1 mM EDTA). Meanwhile, a 20 μl solution containing 0.5 × phosphate buffered saline (Sigma-Aldrich), 10 μl DMSO, 1 mM dibenzylcyclooctyne-amine (Sigma-Aldrich) and 12.5 mM Atto 647N-NHS ester (Sigma-Aldrich) was incubated at 4°C for 1 hour. The DNA sample was split into 30 μl aliquots and 10 μl of the mixture containing the Atto 647N was added to an aliquot. This mixture was incubated at room temperature overnight, before purification by GenElute Bacterial Genomic DNA kit and eluted into 50 μl TE Buffer (10 mM tris, 1 mM EDTA).

### Molecular combing

Molecular combing of DNA was performed based on the procedure described by Deen et al (15). Glass coverslips (Borosilicate Glass No. 1, Thermo Fisher) were cleaned to remove any fluorescent contaminants by incubation in a furnace oven at 450°C for 24 hours. After removing from the furnace and allowing to cool, 30 μl of Zeonex solution (Zeon Chemicals, 1.5% w/v solution Zeonex 330R in chlorobenzene) was deposited onto a coverslip on a spin coater (Ossila) and subsequently spun at 3000 rpm for 90 seconds. Zeonex-coated coverslips were allowed to dry at room temperature overnight and stored in a desiccator.

To perform the molecular combing, 2μl Atto 647N-labelled DNA (2ng/μl in 1×TE) was suspended in 17μl 100mM sodium phosphate buffer (pH 5.7) containing 1μl DMSO. A 1.5μl droplet of this solution was deposited on the surface of the Zeonex-coated coverslip. A clean pipette tip was placed in contact with the droplet and used to drag it, with a velocity of approximately 5mm/min, across the coverslip.

### Fluorescence microscopy

Deposited DNA was imaged on an ASI RAMM microscope, equipped with a Nikon 100x TIRF objective. Illumination was from a 100mW OBIS 640 nm CW laser via a quad-band dichroic mirror (405/488/561/635) and images were collected using an Evolve Delta EM-CCD camera, via a quad-band emission filter (Semrock, 432/515/595/730 nm). Micromanager was used to control the system and scan the sample (16).

### DNA barcode extraction, pairwise alignment and community detection

Software was written in MATLAB (R2016b, The MathWorks, Inc., Natick, Massachusetts, United States of America) for the automated extraction of DNA barcodes from microscopy images, in silico generation of DNA barcodes and the alignment procedures. The computational process is outlined in Figure 1. Full details of the procedure for these processes are given in the Supplementary Information.

**Figure 1.**
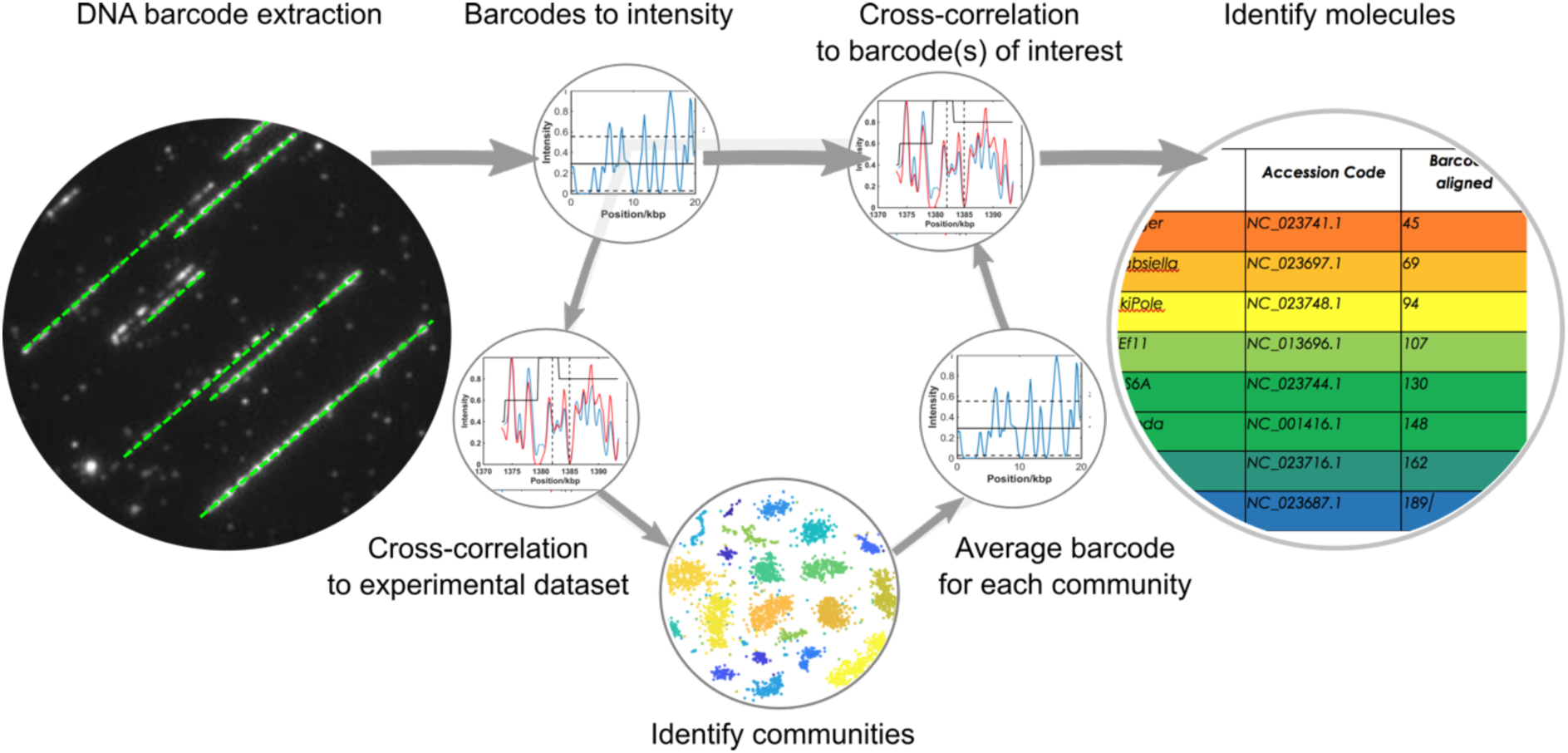
Overview of the computational steps taken to extract DNA barcodes from images and match them to a genome. Intensity profiles for each extracted barcode can either be directly compared to a database of molecules of interest or to every other barcode in the experimental dataset. In the latter approach, communities of similar DNA molecules can be identified and an average barcode for each community determined. Hence a single barcode for each community can be matched to a large database of possible genomes.

## RESULTS AND DISCUSSION

We label DNA using the M.TaqI DNA methyltransferase enzyme to direct the conjugation of fluorophores to sites reading 5’-TCGA-3’ (one site every 256 bp, on average). The M.TaqI enzyme is well-suited to DNA labelling using synthetically-prepared analogues of *S*-adenosyl-L-methionine and we use it here to add reactive azide groups to target adenine bases. We tested a range of conditions for labelling and several DBCO-conjugated organic dyes and developed conditions that give approximately 9 fluorophores for every 10 labelling sites on long, genomic DNA molecules (13, 14). Labelled DNA was separated from the methyltransferase enzyme and reactive dye using a standard silica-based column (for genome purification).

In a typical experiment, we deposit hundreds of gigabases of DNA barcodes from a 1 µL droplet containing 100 pg of DNA(16). A fraction of this, typically a few gigabases, is imaged for analysis. We visualize of the order of 10,000 molecules in 1000 fields of view in ∼20 minutes of imaging (Supplementary Figure S2 shows an example dataset). Software for the automated extraction of the images of DNA molecules from these datasets was developed and is described in the Supporting Information and in Figure S3. Each DNA barcode extracted from the imaging data is stored as a string of integers, where each integer represents the intensity of an individual pixel along the DNA image. Initially, this dataset is filtered to remove molecules that are entwined/aggregated (using an average intensity threshold) and those that are estimated to be shorter than 30 kb in length.

### An *in silico* assessment of the impact of experimental variables on mapping

We generated DNA barcodes (as strings of integers) *in silico* and used these in Monte Carlo simulations in order to understand the impact of a range of experimental variables on our ability to match a DNA barcode to a given DNA sequence. Barcodes were generated from the *E.coli* K-12 genome sequence using the parameters described in Table S1. A summary of the results from these simulations is given in the Supplementary Figures S4 and S5. We find that, for example, for DNA molecules >30 kbp in length, a labelling efficiency of one fluorophore per palindromic target site will allow accurate matching of 90% of the experimental data to the 4.6 Mbp *E.coli* genome, Supplementary Figure S5. The ability to match such short DNA barcodes to a reference genome highlights a significant advantage of using a high density of DNA labelling for the mapping experiment. The preparation of ultra-long DNA molecules is time-consuming and requires significant expertise, yet molecules 30-50 kbp in length can be readily prepared using standard sample preparation kits for genomic DNA extraction and purification.

We also used the barcodes we generated *in silico* to improve our measure of the goodness of fit between a barcode and its reference sequence. These calculations show that an approach relying solely on the normalized cross-correlation between barcodes does not give a sufficiently discriminating measure of the goodness of fit to resolve correctly- and incorrectly- fitted populations of molecules (for the simulated sample of *E.coli* K-12 genome). In order to address this, we introduced an alignment weighting, which is calculated as the mean of three measures of fit quality; the normalized cross correlation, the difference in intensity of the two signals and the difference in the gradients of the two signals. By doing so, we were able to improve significantly the accuracy with which we could resolve correctly-aligned molecules from an incorrectly-aligned population, even with relatively low labelling efficiencies (Figures S6 and S7).

### Pairwise matching- Locating specific genomic regions

In a simple implementation of our mapping approach we make a pairwise comparison between an imaged DNA barcode and a barcode generated *in silico* for a genome or genomic feature of interest. Figure 1 describes this analytical pipeline for matching a DNA barcode to a known reference sequence. Cross-correlation of the signals of thousands of molecules with this (short) reference sequence in this instance takes around ten seconds (standard laptop computer with 16 GB RAM, 3.20 GHz Intel Core i7 processor). Subsequently, an alignment weight is generated and used as a measure of the match quality. Molecules with a weight above a specified threshold are used to generate an average experimental barcode that can be inspected to visually confirm the match to the reference data. This works well in the case where the reference data is a short (say 50-100 kilobase pair) sequence of interest. However, the time taken for matching the experimental data scales linearly with the number (total size) of the genomes in the reference library. Hence, for example, running the sample shown in Figure 2 against a library of 2000 virus genomes would take around 5 hours. The likelihood of a spurious match occurring also increases with the size of the reference library. Supplementary Figure S8(A) shows the result of matching of individual barcodes from an experiment containing the bacteriophage lambda and T7 bacteriophage genomes against a library of twenty virus genomes. Whilst both the lambda and T7 genomes are well represented in the dataset of correctly matched molecules, neither can be reliably identified by simply assigning barcodes to their best match in the reference database. However, an identification can be made with greater than 80% accuracy by using an appropriate weighting threshold (Figure S6, S7 and supporting text) and counting the number of molecules with matches above this threshold for all genomes in the database, Figure S8(B).

**Figure 2.**
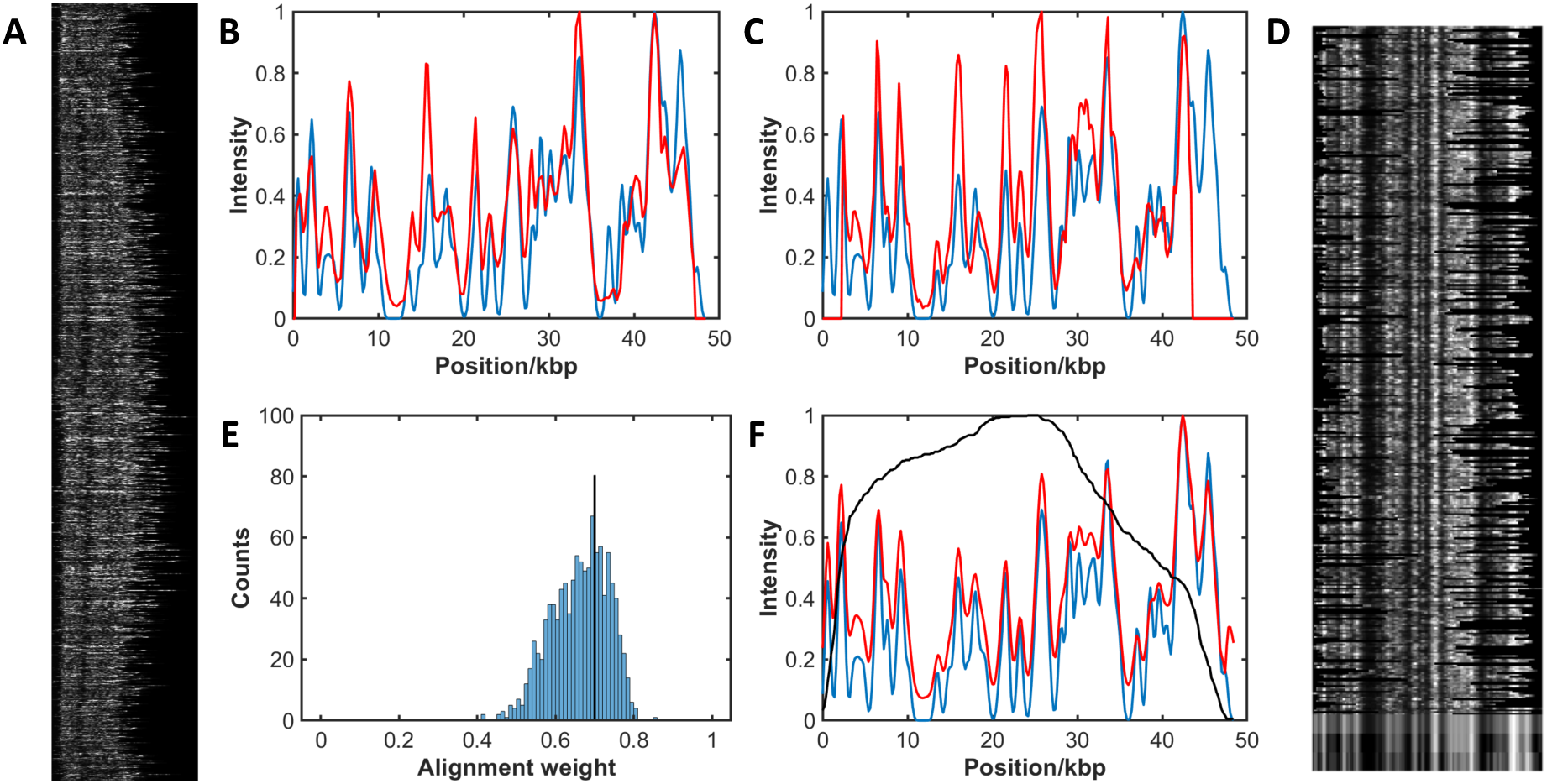
Alignment of pure experimental sample of the bacteriophage lambda genome. DNA was labelled with Atto647N using M.TaqI to direct labelling, combed and imaged. 38,000 candidate DNA barcodes were extracted from the images. A) After filtering the experimental data, 1077 barcodes are identified for further analysis. B-C) The fluorescence intensity profile of each experimental DNA barcode (red) is cross-correlated with a reference molecule (blue) to find the optimal stretch and alignment of the data. D) Selected barcodes (368) with an alignment weight greater than the threshold. At the bottom of the image is shown the mean experimental barcode, the mean with the background removed and the reference barcode (top to bottom). E) Alignment weight for all experimental barcodes is calculated and a threshold (of 0.7 in this case) is applied (black line). F) Plot of mean experimental barcode (red) against the reference barcode (blue), with the number of barcodes (black).

We now extend this approach to a ‘real-world’ example to identify a specific genomic region of interest. We consider a sample of the *E.coli* strain DH10B (TOP10) genome, which has been edited using CRISPR/Cas9. Although the edit in this case is only 64 base pairs in length (∼30 nm on the stretched DNA) and so cannot be directly visualized using standard optical microscopy, molecules matching the region of interest can be identified from their barcodes and we can subsequently search the original dataset for the images of those molecules, Figure 3. Supplementary Figure S8 gives a second example of a search for the *lacZ* gene in this strain of bacteria. Overall, this approach enables interrogation of the genomic imaging dataset for specific features of interest, based on an underlying DNA barcode (sequence). Note that there are several ‘gaps’ in the barcode alignment, where none of the experimental barcodes are aligned to a particular region. On further investigation, we found that this is due to the reference DNA barcode at these locations, which has an inherently low alignment weighting. Hence, even the best matches in these regions have alignment weightings that lie below the cut-off threshold that we have applied (uniformly) across the whole genome. Future work will focus on developing a dynamic threshold for any reference genome (a ‘p-value’) that allows an unbiased estimation of the appropriate threshold for a ‘good’ match at a given locus.

**Figure 3.**
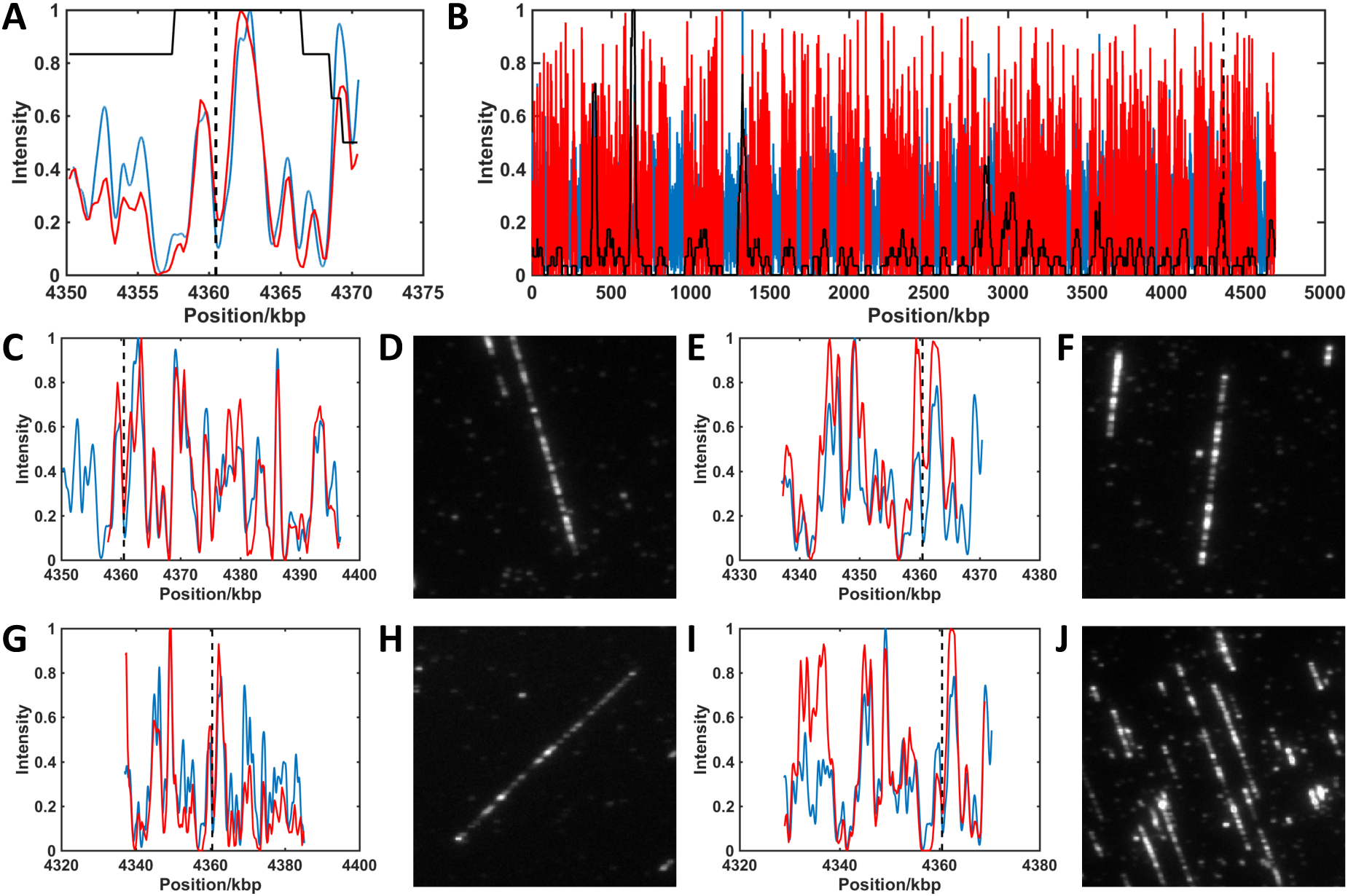
Localisation of barcodes containing a CRISPR/Cas9 edit. Red lines show experimental barcode profiles. Blue lines show reference genome profiles. Black dashed lines show the expected position of the edit on the genome. Values in parenthesis below show alignment weight to reference. A) Consensus barcode generated from all barcodes overlapping with at least 25% of the region of interest. A maximum (minimum) of 35 (18) barcodes (solid black line) contribute to the consensus (0.963). B) Consensus of all barcodes aligned to the genome reference. A maximum (minimum) of 168 (0) barcodes (solid black line) contribute to the consensus across the genome (0.815). C,E,G,I) Single molecule barcodes aligned to region of interest (0.866, 0.864, 0.860, 0.831). D,F,H,J) Raw images of barcodes (shown in C,E,G, and I, respectively) identified as overlapping region of interest.

This analytical approach can be readily extended to strain-type cultured bacteria rapidly. The genomes of three samples, *E. coli* strain DH10B; *E. coli* strain EC958 and *K. pneumoniae* strain Ecl8 were imaged and, using this simple pairwise comparison of DNA barcodes to reference genomes, we were able to successfully strain-type the samples. Around 1000 DNA barcodes from each dataset were matched to reference sequences in a two-step process to identify the species and strain of bacterium against a library of 25 species, (Figure 4).

**Figure 4.**
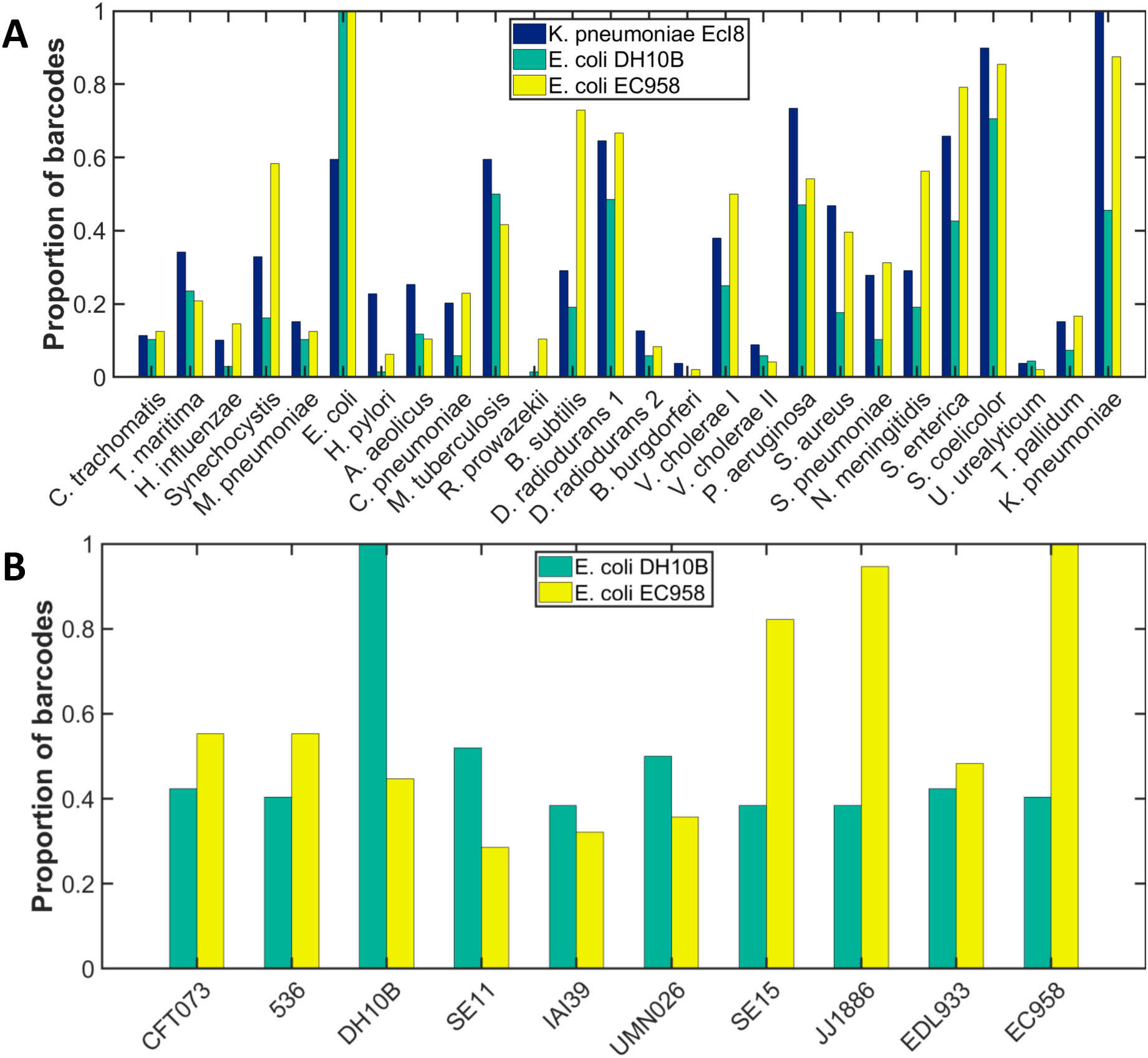
Identification of bacterial DNA by species. Samples of DNA extracted from cultured *K. pneumoniae* strain Ecl8 (blue); *E. coli* strain DH10B (green) or *E. coli* strain EC958 (yellow) were labelled with Atto 647N using methyltransferase-directed labelling (M.TaqI). (A) Each barcode from the sample was assigned to the species to which its alignment yielded the highest alignment weight. In each case, a simple count of the number of barcodes in the sample matching a given reference genome in this way, allows the specie of bacterium to be identified. (B) Similarly, having identified a species, each barcode is now matched against several bacterial strains and then assigned to the strain to which its alignment yielded the highest alignment weight. Both *E.coli* strains are correctly identified. In both plots the bars are scaled such that the most populated field has a value of 1. Accession numbers for the genomes used and the absolute numbers of barcodes assigned to each are given in the Supporting Tables S2 and S3.

### De novo clustering of similar genomic barcodes

The mapping dataset offers a unique way to investigate the genomic composition of a sample without prior knowledge of its content. We sought to take advantage of this by developing an approach to cluster, quantify and identify similar molecules in the dataset. In the mapping data, connections between molecules that are highly similar can be made using the alignment weighting. This allows us to plot individual DNA barcodes in a 2D space, where the proximity between points in the plot, each of which represents an individual barcode, describes their similarity or connectedness. In this way, communities of related barcodes can be identified and these can be used to generate single, consensus barcodes, representative of their community. Such analysis encompasses many thousands of imaged molecules but provides a mechanism for dramatically reducing the size of the dataset and then comparing this reduced representation to a large reference library. For 1000 (∼40 kbp) experimental barcodes, the process of generating these communities of similar barcodes takes approximately one minute.

We validated this approach using a total of 6000 DNA barcodes generated in silico; 4800 of these are from a selection of 20 viral genomes and the remaining 1200 are randomly generated to represent, for example, contamination of the sample or poorly labelled DNA molecules. The analysis groups these molecules into a series of distinct clusters, visualized using a tool for dimensional reduction, t-SNE(17), in Figure 5. The t-SNE plot displays each DNA barcode as a single dot and clusters similar barcodes closely. Around the periphery of the plot is a ring of barcodes that are equally dissimilar to all other barcodes in the dataset (the predominantly grey colouring of the dots indicates that these are almost exclusively the randomly-generated DNA barcodes in the dataset). The consensus barcodes for each detected community were generated and subsequently aligned against a library of 2000 phage genomes, a process that took around 1 hour to complete. Figure 5C and Table S2 summarise the output of these alignments for five shuffles of the dataset. Randomly re-ordering (shuffling) the dataset changes the relative positions of any two molecules in the affinity matrix for that dataset. We use this to ensure that we recover similar communities for each shuffle and, hence, can be confident that the process we use for constructing affinity and adjacency matrices is robust, with respect to the order of the dataset.

**Figure 5.**
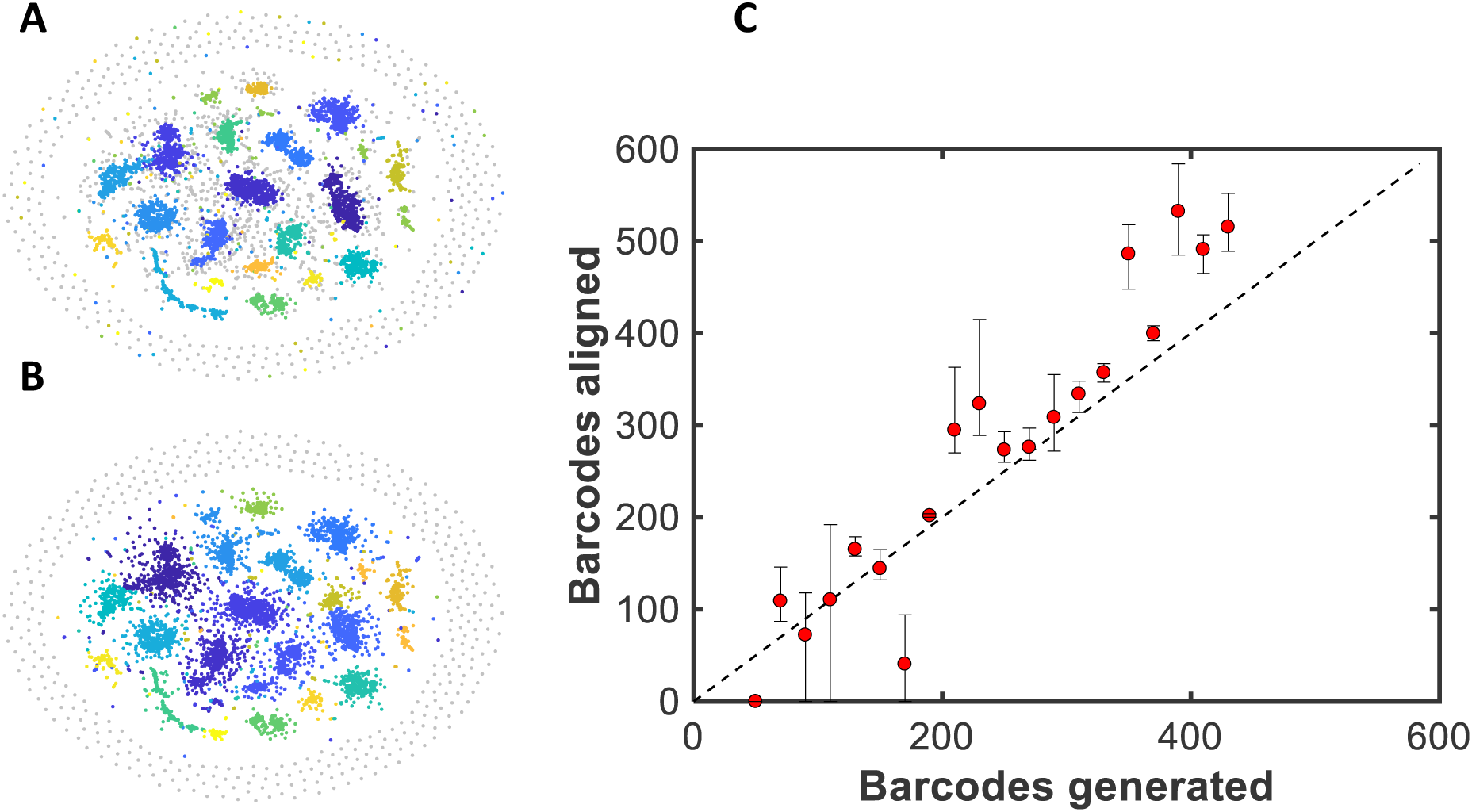
Community detection for barcodes generated in silico. For each of 20 bacteriophage genomes, between 50 and 430 barcodes were generated, with 50% labelling efficiency (coloured dots). 1200 randomly generated barcodes were also added to the simulation (grey dots) A) t-SNE visualisation of the network of communities generated from the synthetic data. Each colour represents a different genome with the grey dots representing randomly-labelled DNA barcodes. B) t-SNE visualization of the communities detected in the dataset. Each colour represents a community of similar DNA barcodes. Note that communities tend to absorb similar random barcodes. As expected, a ring of predominantly random barcodes, which are poorly matched to all other barcodes in the sample, appears on the periphery of the t-SNE visualization. C) Plot showing the mean number of barcodes matched against those generated. Data was shuffled and aligned five times with error bars showing the maximum and minimum number of barcodes assigned to a given genome for the five repeats.

Figure 5 shows that, using data generated *in silico*, 18 of the 20 bacteriophage genomes that were introduced to the mixture can be consistently identified across all five repeats of the analysis. Of the other 1974 genomes in the library, no incorrect (false positive) matches were made. The KBNP1711 genome is not identified in any of the analyses and we attribute this to the inherently low score our alignment weighting gives to good matches with this genome. Other factors, such as absolute and relative barcode numbers, as well as coverage, also play a role in determining the efficacy of the analysis detecting a given genome. Also, note that the impact of the randomly generated barcodes on the analysis limits an absolute, quantitative analysis of the data at this point. Such barcodes represent, for example, a contaminating population of DNA in the dataset or poorly labelled DNA molecules. The detected communities (Figure 5B) invariably expand to absorb a handful of these molecules, such that we generally overestimate the number of barcodes belonging to a given community in the analysis.

To test this approach experimentally, we sought to apply it to identify viral infections of cells. In a first example, we doped a known virus genome (bacteriophage lambda) into a sample of bacterial genomic DNA (*E.coli* ER2566), which was labelled and imaged, as described earlier. We mixed the sample at a ratio of 1:4 (weight for weight virus: bacterial genomic DNA) to mimic the expected copy number- around 20 copies per cell- of such an infection.

Figure 6 shows a clear and unambiguous identification of the phage lambda genome from the mixed sample. In this case, we extracted 5114 barcodes from the imaging dataset, of which 475 were clustered and identified as the lambda phage genome. This process, run against a library of 2000 phage genomes took 47 minutes to run on a standard laptop computer (16 GB RAM, 3.20 GHz Intel Core i7 processor). The t-SNE visualization of the data in Figure 6 is typical of an experimental dataset, where (amongst other factors) labelling efficiency, impurities in the sample and aggregation of DNA barcodes give rise to larger, more diffuse communities than we observed for the simulated dataset. Hence, as expected from the simulation, the ‘lambda’ community contains some barcodes from the *E.coli* genome and we overestimate the number of lambda phage genomes in the sample, attributing almost 1 in 10 of the barcodes to lambda, whereas we introduced approximately 1 in 20 to the mixture. Note that the association of these imperfect/contaminating barcodes with the lambda community has no significant impact on our ability to accurately identify the genome from a library of possibilities, using the consensus barcode of a given community.

**Figure 6.**
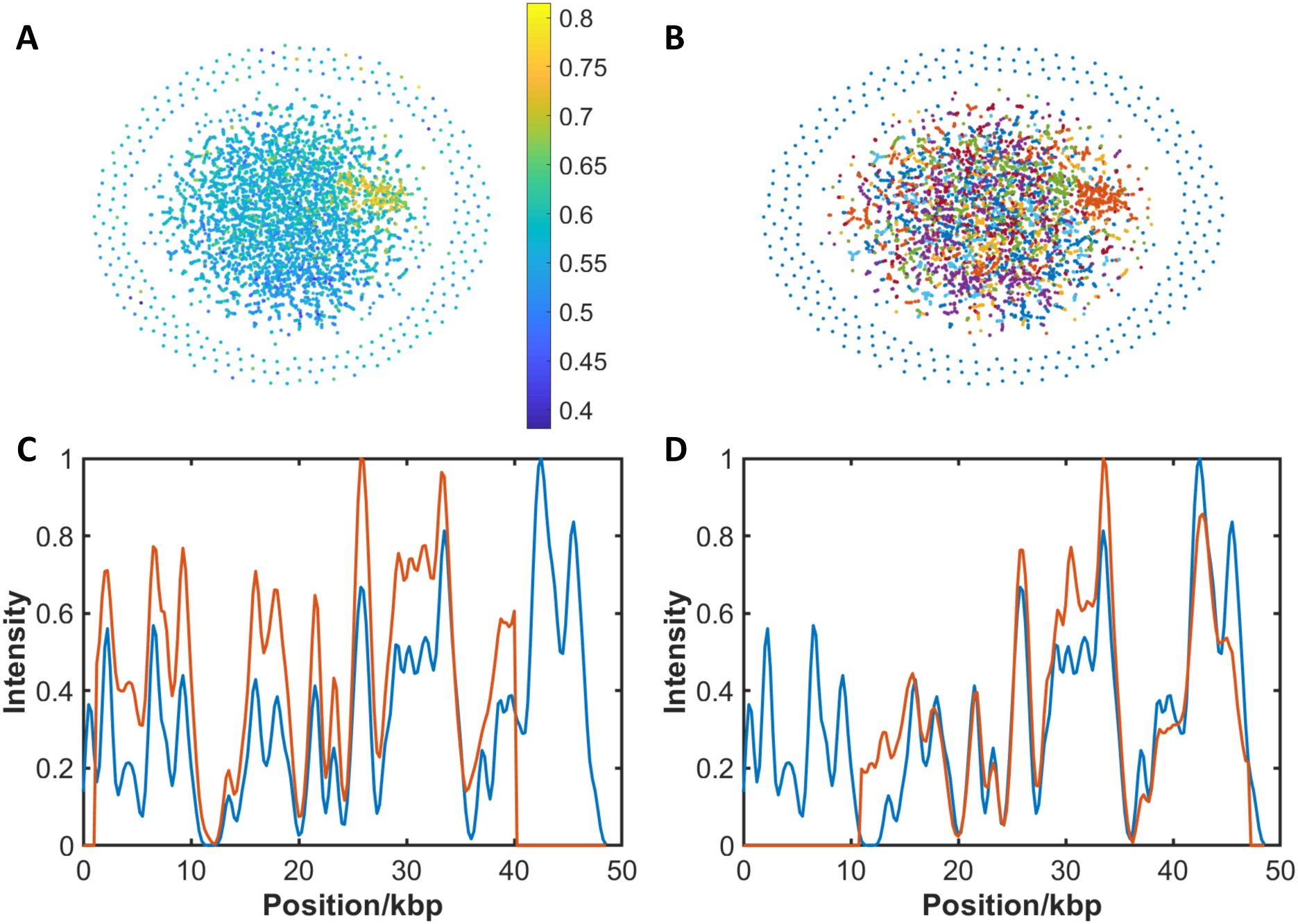
Identification of Lambda bacteriophage DNA in genomic mixture by de novo separation and alignment of experimental barcodes. A) t-SNE visualisation of a network generated from the adjacency matrix. Colour is given by alignment weight to lambda reference genome. B) Community detection. Each colour represents a community that has been detected. 475 of 5114 barcodes are assigned to Lambda phage DNA. C-D) Examples of alignment of consensus barcode generated from clusters.

We extended this model study to investigate an infection of a population of cultured human (HeLa) cells, infected with human adenovirus A (type 12), 72 hours, post-infection. The mixed sample of human and viral DNA was fluorescently labelled, deposited and imaged. The barcodes extracted from these images (1986) were compared to one another, creating an affinity matrix from which 21 communities were identified. Consensus barcodes from these were compared to a database of 128 reference barcodes of vertebrate viruses. A single virus (human adenovirus A) was identified from the sample, in approximately 10 minutes, Figure 7. Supplementary Figure S10 shows exemplars of individual virus DNA barcodes identified in this approach.

**Figure 7.**
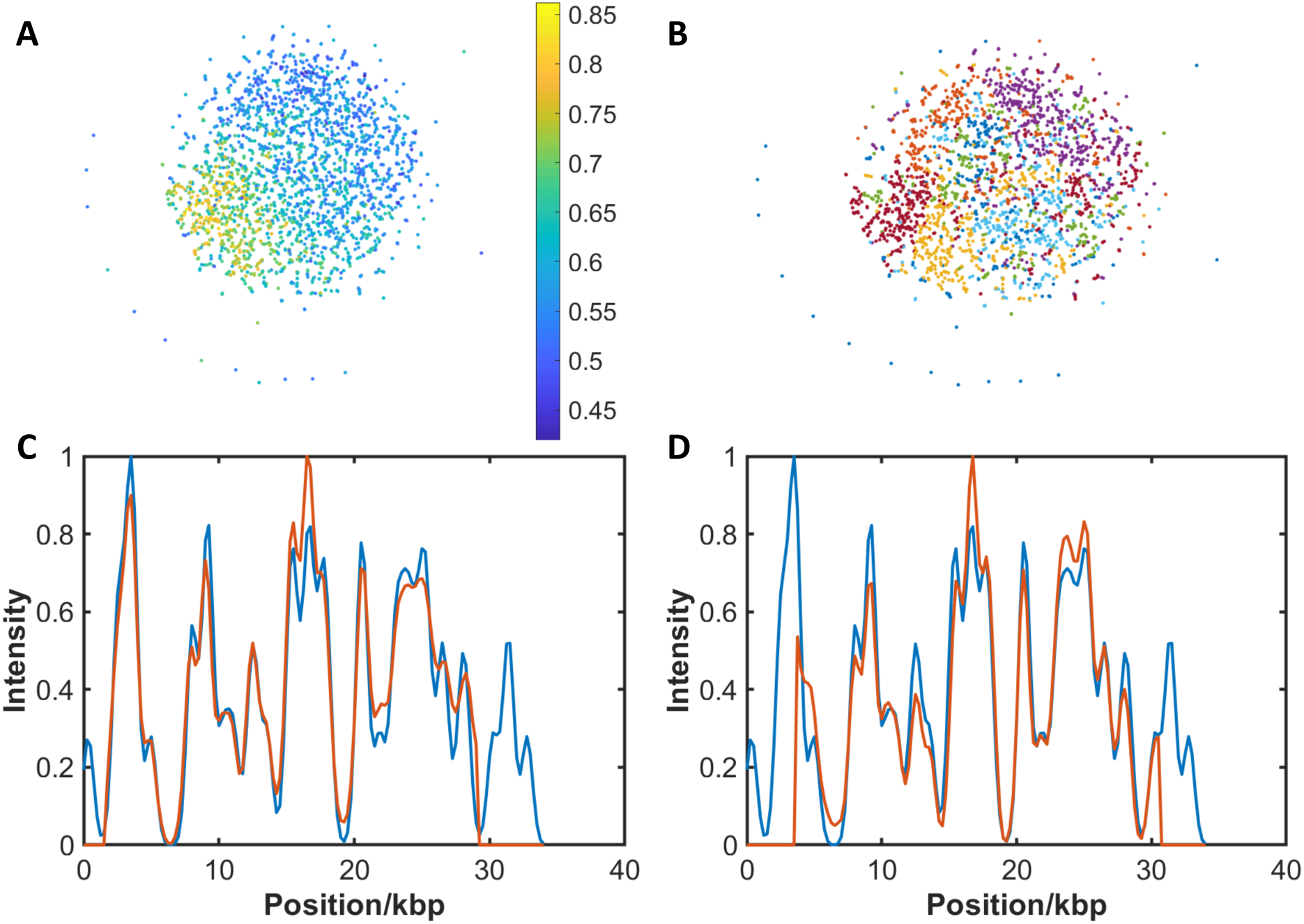
Identification of human adenovirus A DNA sample by separation, de novo alignment and assignment of consensus barcode to reference library. A) t-SNE visualisation of a network generated from the adjacency matrix. Colour is given by alignment weight to adenovirus A reference genome. B) Community detection. Each colour represents a community that has been detected. 669 of 1986 barcodes are assigned to adenovirus A DNA. C-D) Examples of alignment of consensus barcode generated from communities.

In the two above examples, each cell in the sample contains several tens of copies of an invading genome. We sought to explore the limits of our approach by applying this to the identification of (relatively low copy number) plasmids in bacterial cultures. Initial simulations of mixed samples of genomic DNA and plasmid DNA showed promising results, even where we introduce appropriate levels of experimental imperfections to the synthetic dataset. However, initial experiments using this approach have proven unsuccessful in reliably identifying large plasmids in bacterial cultures. This we attribute to their low copy number in bacterial cells. Indeed, if we reduce the number of copies of a plasmid to less than five per cell, simulations show that it is (unsurprisingly) difficult to identify a community of barcodes that can be mapped to a specific plasmid. However, since the sequence of interest is known in this case, a simple pairwise alignment of the dataset to a reference (plasmid) barcode can be used and has yielded a handful of candidate barcodes with high alignment weighting to the resistance plasmid (Figures S11-S13). Further investigation is required to confirm the identity of these molecules, yet this preliminary result suggests that the identification of a small number of molecules of interest from a large sample is possible. Such an approach could be twinned with super-resolution imaging in the future to better resolve the barcodes of such a subset of candidate molecules.

## CONCLUSION

We have provided a basis for the use of DNA barcodes as a tool for visualization of the sequence of whole genomes. We have collected multiple datasets containing gigabases of DNA, each of which is of the order of 120 megabytes in size. We have shown that this data can be searched for a genomic region of interest, used to search a library of genomes for matching organisms and that similar molecules within the dataset can be identified and clustered to identify viral infections against the background of host genomic material. In the future, work will focus on exploiting the strength of these optical measurements to allow multiplexed studies of both sequences and ‘events’. Indeed, we believe that this unique view of the genome will enable a new perspective on events and processes, such as replication, drug or protein binding or genomic editing across whole genomes, at the single-molecule level.

## AVAILABILITY

The datasets used for analysis in this study and an annotated version of the Matlab code we used to extract, process and analyse DNA barcodes are available at edata.bham.ac.uk, DOI: 10.25500/eData.bham.00000255.

## ACKNOWLEDGEMENT

We would like to gratefully acknowledge Dr Francisco Fernandez-Trillo for invaluable support regarding the synthesis of AdoHcy-azide, Dr Roger Grand for supplying DNA extracted from HeLa cells, infected with the Human adenovirus; Dr Michelle Buckner Prof. Laura Piddock for supplying DNA extracted from bacterial cultures. We would especially like to thank Anna Dumitriu and her collaborator Dr Sarah Goldberg (MRG-Grammar/Technion) for sharing with us her ‘Make Do and Mend’ strain of *E. coli* and for bringing her inspirational art/science approach to our lab.

